# Uncovering disease progression trajectories across multiple timescales with hybrid ODE-mixture models

**DOI:** 10.64898/2026.10.29.762792

**Authors:** Max de Rooij, Jesse D. Quadt, Natal A.W. van Riel, Shauna D. O’Donovan

**Affiliations:** Dept. of Biomedical Engineering, Eindhoven University of Technology, Eindhoven, the Netherlands; Eindhoven Artificial Intelligence Systems Institute (EAISI), Eindhoven University of Technology, Eindhoven, the Netherlands

## Abstract

Biological systems are capable of evolving across multiple timescales, yet most modeling approaches capture either short-term physiological responses or long-term disease progression. Additionally, methods that combine these time scales either struggle with inter-individual variation, or require large volumes of data to work. We present Latent Class Parameter Trajectory Analysis (LCPTA), a hybrid framework that combines mechanistic ordinary differential equation (ODE) modeling of short-term dynamics with latent class trajectory modeling of long-term parameter evolution. To support this approach, we developed TemporalMixtureModels.jl, a Julia package for univariate and multivariate temporal mixture modeling. We evaluated LCPTA in two synthetic longitudinal studies, covering a simulation of a heterogeneous population developing hypertension, as well as a synthetic longitudinal study of 75 virtual individuals followed over 30 years, representing muscle insulin resistance, liver insulin resistance, and healthy trajectories. We compared LCPTA with a conditional neural ODE and a conditional neural network for reconstruction of long-term parameter trajectories and subgroup separation. In both studies LCPTA recovered relevant subgroup structure and trajectory patterns. In the cardiovascular study, the neural approaches were able to distinguish inter-individual differences, but in the insulin sensitivity study, neural approaches failed to represent parameter trajectories faithfully, primarily separating individuals along a single dominant parameter and showing weaker multivariate trajectory reconstruction. In additional experiments with sparse and irregularly sampled unseen individuals, LCPTA maintained strong classification performance. We show that LCPTA is an efficient and interpretable strategy for multiscale modeling of heterogeneous metabolic disease progression, with potential utility for data-efficient digital twin applications.

**Author summary:** Development and progression of disease is a complex process, which can be understood through the slow dysregulation of a biological system over time. Typically, biological systems operate on a different time scale than the dysregulation occurs. In order to model disease progression, we therefore require mathematical models that operate across these different time scales. In this work, we present latent class parameter trajectory analysis (LCPTA) as a methodology to combine mathematical models across multiple time scales. This methodology works by combining ordinary differential equation (ODE) models on their intrinsic time scales with latent class trajectory analysis (LCTA), to describe the gradual change of ODE model parameters on the time scale of dysregulation. We present two simulation studies of hypertension development and insulin resistance development, demonstrating the ability of LCPTA to reveal relevant different groups in disease progression. We compared LCPTA to approaches presented in previous work involving neural network modelling of the dysregulation time scale, and show that LCPTA matches neural network performance on univariate parameter trajectories, and strongly improves over neural network models on multivariate parameter trajectory analysis. To support the use of LCPTA in future work, we have built and released the Julia package TemporalMixtureModels.jl.

## Introduction

The development and progression of human disease is inherently complex, characterised by the gradual dysregulation of a biological system over time. These biological systems typically consist of many interconnected and hierarchical components that operate over diverse time scales. [1] Moreover, biological systems adapt to changing environments. [2] The short-term dynamic responses to external stimuli observed over minutes to hours, such as the increase in heart rate in response to stress or exercise or the glycaemic response to the ingestion of a meal, give an indication of the body’s fitness or flexibility, often dubbed resilience. Over the longer-term the ability of the biological system to adapt to these external perturbations can also deteriorate, due to age or disease development. Therefore, in recent years, resilience has become central focus for defining health. [3] Resilience has emerged as a central principle in multiple lifestyle-related diseases including type-2 diabetes, where we first see processes regulating glycaemic control gradually become dysregulated through progressive systemic insulin resistance [4], and cardiovascular disease where chronic stress induces increased sympathetic nervous system activity, leading to hypertension. [5]

For both the glucose response, as well as the cardiac cycle, several mathematical models have been developed to study these short-term adaptations. By fitting these models to measured data, we can quantify physical or biological parameters of the system. For example, for models of glucose excursions, parameters are often estimated through metabolic challenges, such as the oral glucose tolerance test (OGTT), where a solution of 75g of glucose is ingested, or Mixed Meal Tolerance Test (MMTT) where a combination of fats, protein, and carbohydrates is consumed. [6–8] By estimating the kinetic parameters of these mathematical models of the glucose-insulin system, interpretable features of an individual’s metabolic resilience can be obtained. [9] These features obtained from a short-term perturbation provide us with information on the current glycaemic resilience status.

However, time scales at which disease progresses are often different from the time scales described in mathematical models. For example, while the plasma excursion of glucose and insulin during an oral glucose challenge occur over minutes to hours, the development of lifestyle-related diseases, such as diabetes type 2, happens over decades. Building on this principle, the values of the model parameters representing a metabolic challenge response may change over the longer time scale indicating disease progression, such as insulin resistance, metabolic-associated steatotic liver disease, or diabetes type 2. [10]

In addition, strong heterogeneity exists in diabetes development trajectories [11]. Mathematical modelling of the long-term change in these model parameters would enable the disentangling of these heterogeneous trajectories. [12] In order to keep these models computationally tractable, the long-term change in parameter values is separated from the short-term parameter values [13, 14]. While modeling of the development of diabetes over the long term has also been performed [15], models that simultaneously describe the evolution of short time scale metabolic responses over the long time scale of disease progression is less common [14, 16].

In order to develop digital twins that can realistically forecast the future behaviors of the biological system, combining both time scales is necessary for consistent personalization over repeated measurements [17].

In previous work, Tiemann et al. [18] have proposed decoupling both time scales by initially estimating kinetic parameters for the mathematical model on the short time scale independently, and subsequently analyzing the evolution of the kinetic parameters over the consecutive measurement instances on the long time scale. This method, labelled Analysis of Dynamic Adaptations in Parameter Trajectories (ADAPT), uses multiple sampled splines through the estimated parameters at each long time scale timepoint to account for possible uncertainty. [13] The key to ADAPT is that the underlying biological process, as characterized by the short-term dynamic model, remains the same while the rates of individual reactions, controlled by the model’s kinetic parameters, may change over time, reflecting a loss in function. A challenge with ADAPT, however, is that it cannot detect structural heterogeneity in parameter trajectories. Furthermore, the use of splines complicates the integration these long-term parameter trajectories in a digital twin application to predict disease trajectories for a new individual.

Building on the work of Tiemann, Regazzoni et al. [14] proposed to solve these challenges by modelling the parameter trajectories using a neural ODE with a latent conditional variable. While, their results showed the neural ODE could recover underlying heterogeneity in a synthetic data set with regularly sampled measurements; in reality longitudinal clinical measurements often have both infrequent and irregular sampling intervals, which complicates full individualization of their neural ODE. Given the long duration of disease development, there also may be limited longitudinal data available. Moreover, heterogeneity in disease development is often complex, potentially containing multiple distinct trajectories, for example an individual with overweight or obesity may go on to develop MASLD, T2DM, cardiovascular diseases or a combination of all three. [11]

To take advantage of potentially shared patterns in long term parameters, while coping with sample sparsity challenges related to long time scale data, latent class trajectory analysis (LCTA) can be used. [19] In this approach, individual trajectories are grouped into a mixture containing a predefined number of clusters. Each cluster is characterized by a set of model parameters, producing a specific model outcome. For unseen individuals, membership probabilities can be computed for each cluster by analyzing the likelihood values for this individual given each model outcome in the mixture. In this way, the model can be used to analyse data for previously unseen individuals. Other names for this modelling approach include latent profile analysis (LPA) [20] or latent class growth analysis [21]. This technique has previously been successfully applied to identify subgroups based on plasma glucose excursions during an OGTT. [22]

In this work, we propose a new modelling-paradigm, labelled Latent Class Parameter Trajectory Analysis (LCPTA), combining LCTA to capture long-term deviations in kinetic parameters with an ODE model of the short-term dynamics, producing a hybrid-framework that can recover distinct subgroups of disease progression from repeated measurements in an unsupervised way. To achieve this, we have developed the `TemporalMixtureModels.jl` package to fit univariate and multivariate mixture models of polynomial regressions on time series data. We showcase the use of the `TemporalMixtureModels.jl` package to efficiently perform LCPTA using multivariate polynomial mixture models on a long time scale to uncover distinct patterns in the deviation of OGTT response, captured by fitting the Eindhoven Diabetes Education Simulator [6], during the development of tissue specific insulin resistance. This concept is illustrated in Fig 1. We use a virtual population of distinct disease development trajectories to test LCPTA’s ability to distinguish different patterns in ODE model parameters. We compare the performance of LCPTA with the neural ODE approach presented in Regazzoni et al. and a regular conditional neural network. Finally, we investigate the performance of inference of disease development class using the trained LCPTA model on an unseen virtual population with sparse and irregularly sampled timepoints.

**Fig 1.**
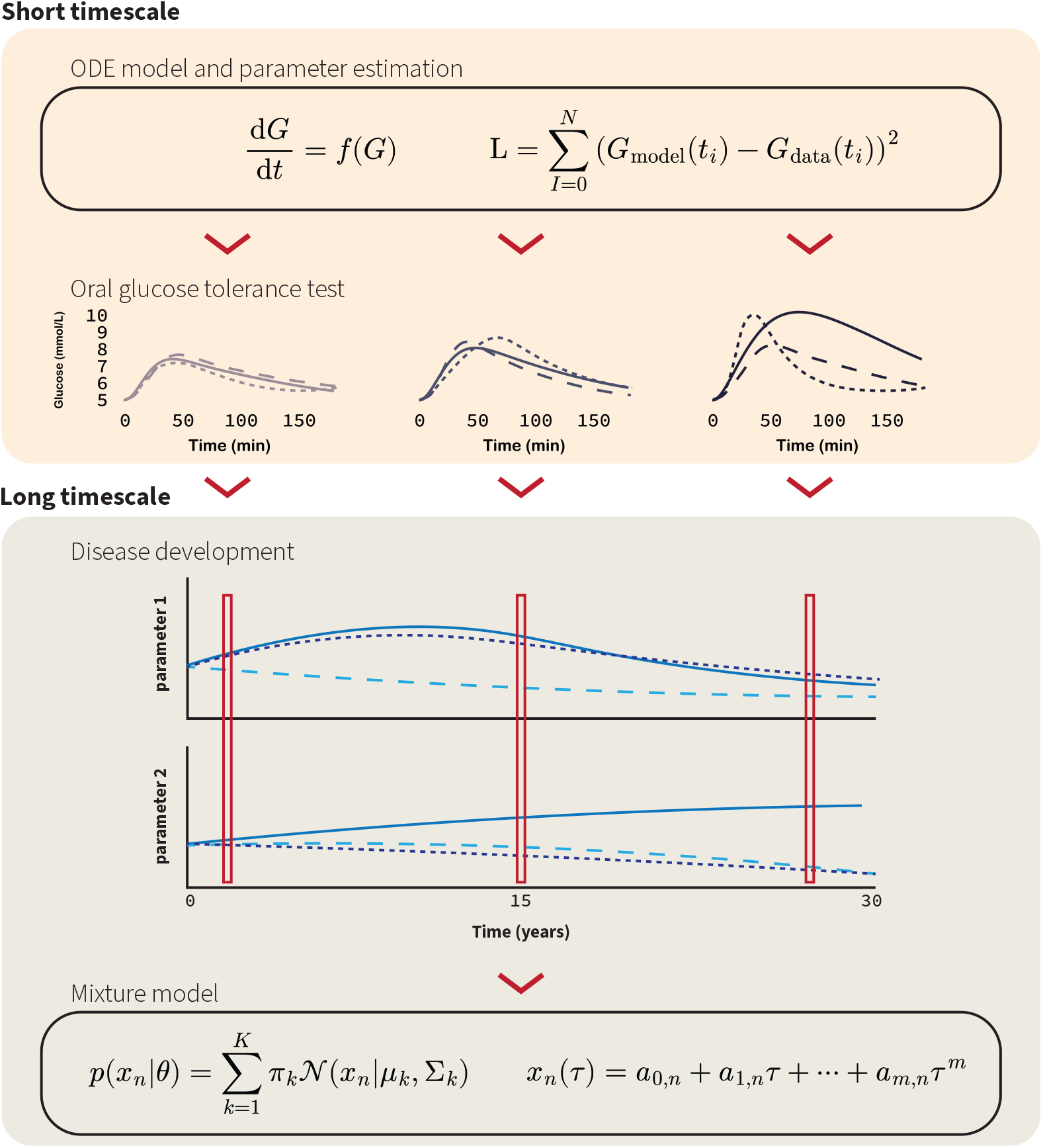
Illustration of the separation of timescales in modelling oral glucose tolerance test glucose excursions and their change over the years. On the short timescale of minutes to hours, we consider the glucose and insulin excursions after an oral glucose tolerance test. These responses have been extensively modelled in previous work, and through parameter estimation with an ordinary differential equation (ODE) model, we can obtain an interpretable set of parameters that describe the glycemic status of an individual. The evolution of these parameters over a long timescale (years) can then be modelled using a mixture of polynomials, effectively grouping similar patterns of disease trajectories.

## Materials and methods

### Generation of Synthetic Data

In order to test the performance of LCPTA, we generated two synthetic longitudinal data sets. The main dataset presented in this work is generated by simulating glucose and insulin excursions for 2 hours after an oral glucose tolerance test (75 g glucose) in 75 virtual individuals, repeated every six years over a 30 year time span. These 75 virtual individuals were split into three groups of 25 individuals representing individuals developing primarily muscle insulin resistance (MIR), primarily liver insulin resistance (LIR), and no insulin resistance (Healthy).

Additionally, we reproduced the synthetic data generated by Regazzoni et al. [14], simulating the development of hypertension using a model of the cardiovascular system. Full details on this dataset can be found in S5 Text.

#### Oral Glucose Tolerance Test

Oral glucose tolerance test data was simulated using the Eindhoven Diabetes Education Simulator (EDES) model, which is an ODE model to simulate postprandial glucose and insulin for people with diverse glucose tolerances. [6] (S1 Fig). Model equations and parameters are included in S2 Text. Data was sampled every 30 minutes for a simulated time of 120 minutes. Normally distributed multiplicative noise was added to simulated data, with a mean of zero, and a standard deviation of 0.05. All progression parameters governing the parameter trajectories are included in S3 Table.

#### Muscle insulin resistance

To simulate the developement of muscle insulin resistance over the long time-scale, parameters describing insulin-dependent glucose uptake (*k*_5_) and transport rate from plasma to insterstitial insulin (*k*_9_) were prescribed to decrease exponentially over time in years (*τ* ) as

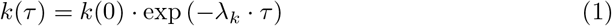

Concordantly, basal glucose and insulin (*G*_*b*_ and *I*_*b*_) were increased over time according to

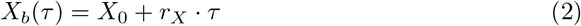

For each individual, progression parameters 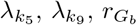, and 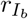 were randomly sampled from population distributions. Additional variability was generated by sampling *k*_3_, *k*_4_ and *G*_*liv,b*_ from random population distributions for each individual and each time point *τ* .

#### Liver insulin resistance

For liver insulin resistance, parameters *k*_3_ and *k*_4_ were selected to decrease according to Eq 1. Parameters *k*_3_ and *k*_4_ reflect the reduction of gluconeogenesis in response to elevated glucose and insulin levels, respectively. As liver insulin resistance is more complicated to accurately simulate using the model, parameters *k*_1_, *k*_6_, *G*_*liv,b*_, *G*_*b*_, and *I*_*b*_ were selected to increase through Eq 2. Furthermore, parameter *k*_5_ was selected to slowly decrease according to Eq 1.

#### Healthy trajectories

For simulated healthy trajectories, natural decline of insulin resistance is simulated by replicating the MIR progressions, but strongly reducing the progression parameters for *k*_5_, *G*_*b*_ and *I*_*b*_, and keeping parameter *k*_9_ constant.

### Estimation of Short-Term Parameter Values

For the model fitting procedure we will follow the parameter estimation strategy that has been developed and used in previous work by Erdős et al. [23], where it was shown that estimating *k*_1_, *k*_5_, and *k*_6_ captures most of the natural variation in OGTT response [23] and is also identifiable. Additionally, we estimate residual standard deviations for glucose (*σ*_*G*_) and insulin (*σ*_*I*_ ). All parameters were independently estimated for each individual and long time scale sample using maximum likelihood estimation.

For individual *i*, at long timescale step *τ*, for glucose data **G**_*i*_ and insulin data **I**_*i*_, parameters *θ*_*i*_ were estimated by minimizing

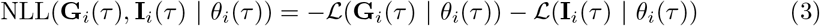

Where ℒ is defined as:

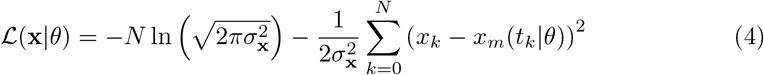

With *N* depicting the amount of data points in **x**, *x*_*m*_ indicates the model prediction of *x*, and *t*_*k*_ indicates the *k*^th^ timepoint in *N* .

### Neural Ordinary Differential Equation to Model the Long-Term Dynamics

To recover the trajectories of the short-term parameter values over the long-time scale, we initially implemented a neural ordinary differential equation (neural ODE) approach proposed by Regazzoni et al. [14], including a conditional latent variable *c*_*i*_ as an additional input to the neural network. In this model, the parameter progressions are described as

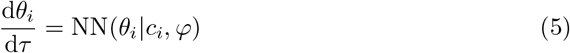

Neural network parameters *φ* are shared between simulated individuals, while the conditional latent variable *c*_*i*_ is estimated per individual to represent inter-individual variability.

Parameters were estimated by minimizing the sum of squared errors between the model and the data for all individuals at once. Optimization was carried out in two steps, starting with the Adam optimizer [24] for 1000 iterations and following with the BFGS optimizer until convergence was detected, or until BFGS ran for 1000 iterations.

#### Conditional Neural Network

For the conditional neural network, we modelled the trajectory as

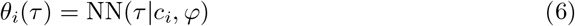

Similarly to the neural ODE approach, the latent conditional parameters were estimated on a individual level, while the neural network parameters were estimated on a population level.

#### Polynomial Mixture Model of Long-Term Dynamics

Here, mixture models are used to model the trajectories of the short-term parameter values over the long-time scale. To fit a polynomial mixture model to the data, we have developed the `TemporalMixtureModels.jl` package. In this package, we have defined the mixture model by default as a polynomial regression function. However, we have deliberately built a customization feature such that other models could be implemented as mixture components.

In this case, each model parameter that was estimated was an outcome variable. The underlying mixture component was then defined as a multivariate polynomial regression with independent outcomes. The model was fit to the data using the implemented expectation maximization (EM) algorithm. To initialize the model parameters (*p*_*k*_) for each component *k*, each individual *i* is randomly assigned to one of the mixture components, and each component regression model was fit to the measurements *D* of its assigned individuals. The EM algorithm then consists of two steps that repeat until convergence or a set maximum number of iterations.

In the first (E) step, each individual *i* is assigned to a mixture component by computing their responsibilities *γ*_*ik*_ for each component *k* according to:]

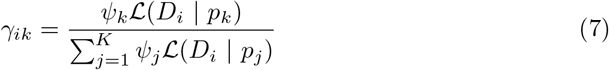

Where *ψ*_*k*_ is the mixture weight for component *k, K* is the number of components, and ℒ (*D*_*i*_ |*p*_*k*_) is the component likelihood for individual *i* and component *k*.

The model parameters of each component are then updated in the second (M) step through a weighted least squares fit, using the responsibilities of each component as weights.

When the change in likelihood value of the model in the current iteration compared to the previous iteration was below a certain tolerance threshold, set here at 10^*−*6^, the algorithm was terminated for convergence reasons.

As mixture models may be sensitive to initial clustering, we repeated every mixture model fit five times using different random initializations, selecting the model with the largest final likelihood.

To select the number of components and the polynomial order of the underlying model, we used 10-fold cross-validation. We varied the number of components between 2 and 5, and the polynomial order between 1 and 4, and used the Bayesian Information Criterion (BIC) to select a set of hyperparameters that balance model accuracy and complexity.

The developed `TemporalMixtureModels.jl` package has been made available through the Julia standard registry and its documentation is available from https://computational-biology-tue.github.io/TemporalMixtureModels.jl/stable/.

## Results

In Fig 2 panels a and b, a visual representation of repeated OGTT responses generated for three virtual individuals in the MIR, LIR, and healthy groups. The difference in glucose and insulin shapes in the simulated OGTTs is clearly visible, where the MIR individual (blue) develops a broad peak, with a slow decline in postprandial glucose, in line with the definition of poor muscle insulin sensitivity proposed by Abdul-Ghani et al. [25], the LIR individual shows a sharp and intense peak of glucose, reflective of incomplete supression of endogenous glucose production in the liver in the early phase of the glucose response (first 30 minutes), which recovers quickly towards fasting values. The healthy progression shows the lowest variability in the glucose and insulin curves. In panels c through e, the trends in the values of k1, k5, k6, the kinetic parameters governing the short-term dynamics re-estimated from the simulated OGTTs using the EDES, model over the 30-year duration of the long-term simulation, are depicted. The MIR individual here (blue) is characterized by a strong dip in *k*_5_, representing the loss of overall peripheral insulin sensitivity. In the LIR individual (red) the dip in *k*_5_ is lower, compared to the MIR individual, but this individual also shows an increase in *k*_1_ and *k*_6_. The heatlhy individual (yellow) has been constructed to show relatively stable parameter values over the 30-year simulation.

**Fig 2.**
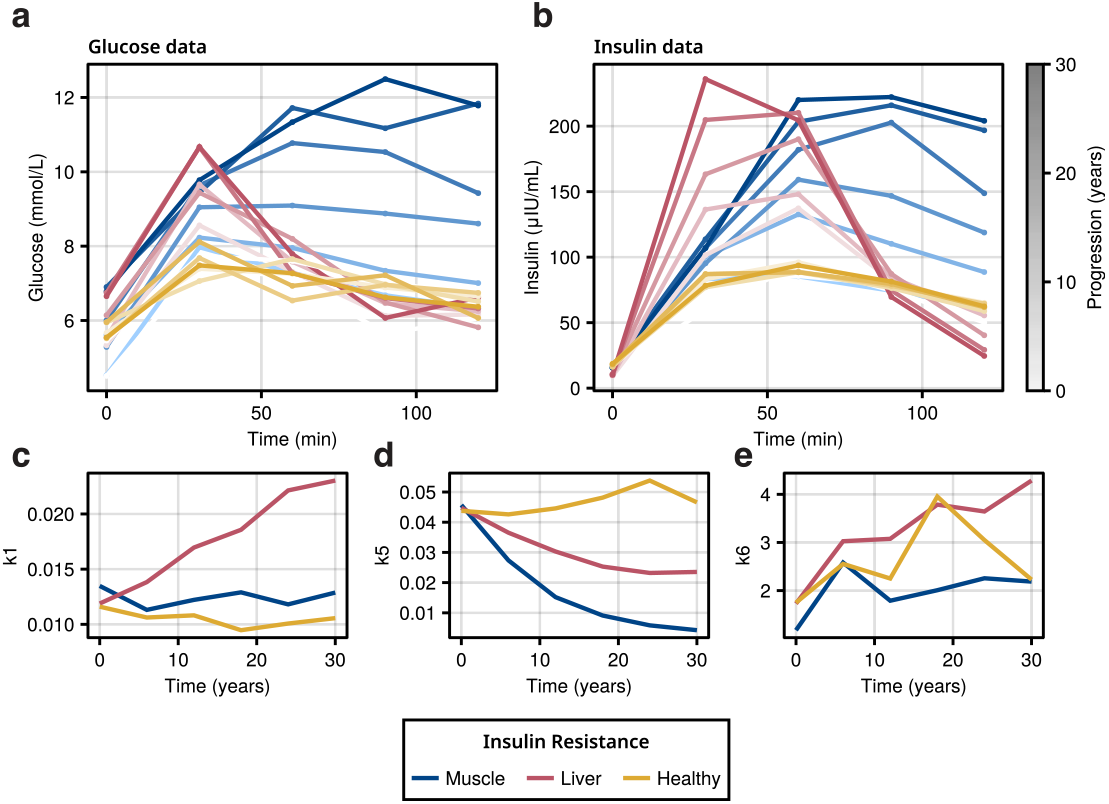
Example of model simulations. A-B: Progression of glucose (A) and insulin (B) measurements during a simulated OGTT for a virtual individual in the muscle insulin resistance (MIR) (blue), liver insulin resistance (LIR) (red) and healthy (yellow) group. The darkness of the color indicates the progression over years on the slow timescale. C-E: Change of estimated model parameter values *k*_1_ (C), *k*_5_ (D) and *k*_6_ (E) over the long time scale for a virtual individual in the muscle insulin resistance (blue), liver insulin resistance (red) and healthy (yellow) group.

### Neural ODE model mainly separates based on a single parameter

We initially tested the neural ODE approach in the simulated windkessel model, replicating the study by Regazzoni et al. [14] We defined a neural ODE containing a conditional parameter *c* to capture the underlying variability. Fig. 3a illustrates the architecture of a neural ODE, where the derivative of the states is estimated by a neural network.

**Fig 3.**
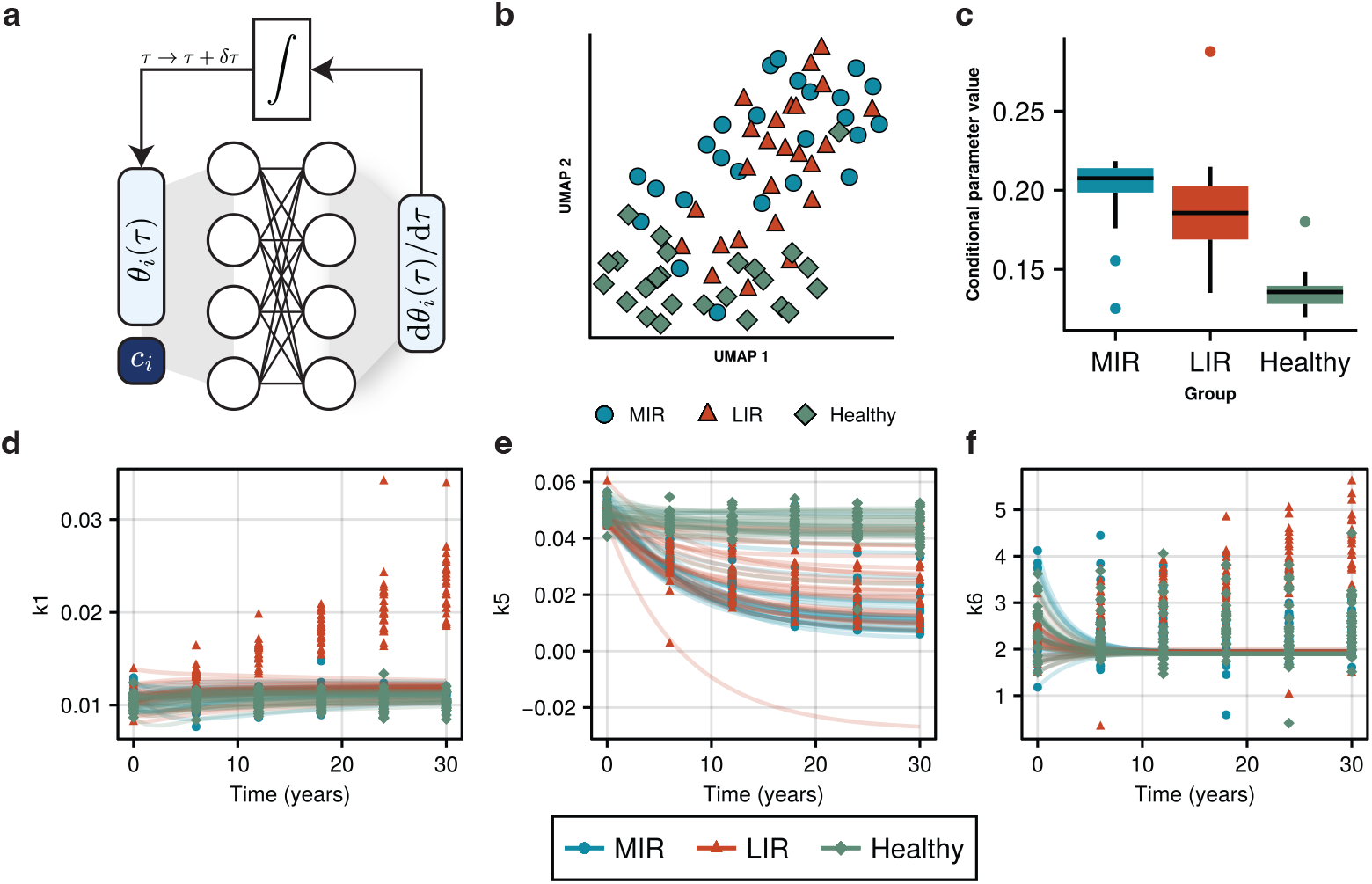
Neural ODE reconstruction of parameter trajectories. Results of the reconstruction of the parameter trajectories of the MIR, LIR, and healthy subgroups using a neural ODE. A: Illustration of the structure of the conditional neural ODE, where a neural network is used to describe the change in parameter values over time (*τ* ). The conditional parameter *c*_*i*_ is constant over time and allows for modelling specific interindividual differences. B: UMAP plot of the trained conditional parameter across 25 optimization runs with different initial parameters, colored by subgroup (blue circle: MIR, orange triangle: LIR, green diamond: healthy). C: Representative boxplot of the conditional parameter values across different groups for a single optimization run. D-F: Reconstructions of the parameter trajectories (lines) for the parameters *k*_1_, *k*_5_, and *k*_6_, respectively. Lines and ground-truth parameter values are colored by subgroup.

In the windkessel model, the underlying variability was encoded in the simulation using a parameter *α*. We ran the optimization procedure 25 times using random initial values for parameters and show that for each of the optimization runs, *c* is strongly correlated with *α* (S8 Fig). Additionally, the model shows a strong agreement with the observed data (S9 Fig).

Next, we used the same neural ODE approach for reconstructing the parameter trajectories in the insulin resistance development simulation.

As conditional parameter values cannot be directly compared between initializations, we assessed whether separate groups could be identified by performing a Uniform Manifold Approximation and Projection (UMAP) on the resulting conditional parameter values across all 25 initializations (Fig. 3b). In the projection, healthy groups are somewhat clustered together, away from the MIR and LIR groups, while MIR and LIR seem to remain close together. A representative distribution of conditional parameters from one of the initializations is depicted in Fig. 3c, showing clear separation between the healthy and the MIR and LIR groups. For the same model as in Fig. 3c, we used the estimated parameters to reconstruct the three parameter trajectories. The parameter profiles (Fig. 3d-f) clearly show a strong focus on *k*_5_, where most of the separation between groups is concentrated, while the other model-predicted parameter profiles show limited distinction between groups and do not follow the trends encoded in the simulated data.

Alternatively, we investigated the difference between a conditional neural ODE and a time-dependent conditional neural network. The conditional neural network aimed at learning the actual temporal parameter trajectories instead of their derivatives. An illustration of the conditional neural network is depicted in S4 Figa.

Similarly to the neural ODE, we used a UMAP to determine whether any of the initializations could capture the differences between the MIR, LIR, and healthy groups.

From the UMAP, we observe that the conditional neural network separates the healthy group from the MIR and LIR groups, in a similar way as the neural ODE and we observe some separation between MIR and LIR. (S4 Figb) Taking a representative model and inspecting the conditional parameter distributions between different groups similarly shows a strong separation of the healthy group and a mild separation of the MIR and LIR groups (S4 Figc).

Subsequent reconstructions of the parameter trajectories, shown in S4 Figd-f, however, reveal that the conditional neural network does not accurately represent the average trajectories nor the variability between groups.

### Polynomial mixture models separate underlying groups

The polynomial mixture model consisted of *k* polynomial regressions, with *τ* as an independent variable. The model is illustrated in Fig. 4a.

**Fig 4.**
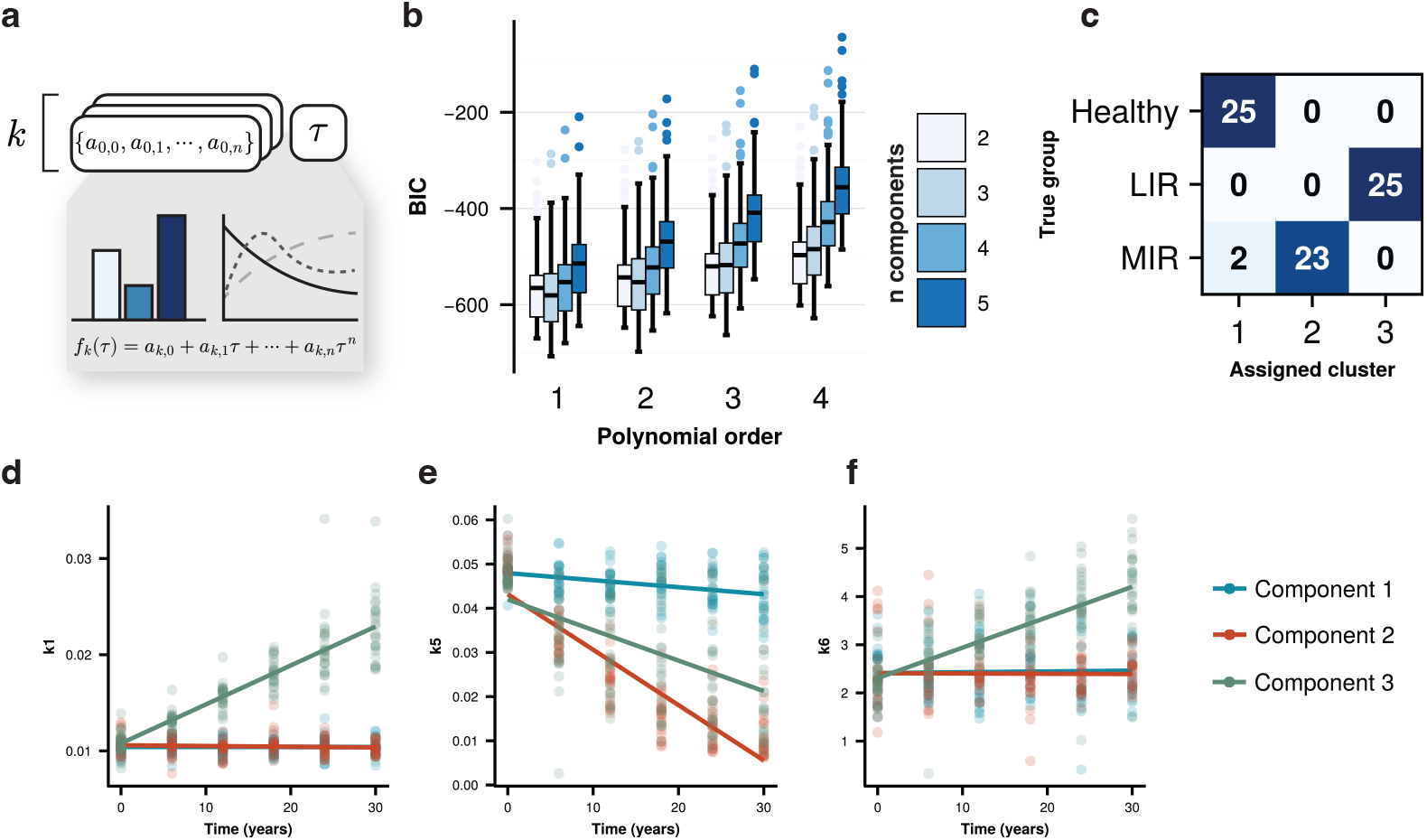
Mixture model reconstruction of parameter trajectories. Results of using a polynomial mixture model to reconstruct the parameter trajectories in the simulated data. A: Illustration of the mixture model as *k* sets of model parameters that each describe a function over time. Each outcome is a weighted combination of outcomes of all functions in the mixture, where the weights change per individual. B: Cross-validation to select the hyperparameters for the polynomial mixture model. The horizontal axis shows the polynomial order of the underlying component models, while the colors indicate the number of unique components *k* in the data. The boxes indicate the Bayesian information criterion (BIC) on the test sets within the cross-validation experiments. C: Confusion matrix comparing the assigned labels with clusters assigned by the mixture model. The total number of individuals per group is 25. D-F: Reconstructions of the parameter trajectories (lines) for the parameters *k*_1_, *k*_5_, and *k*_6_, respectively. Lines and markers are colored by assigned cluster.

To determine the hyperparameters for the polynomial mixture models, we initially performed 10-fold cross-validation on the data, using the Bayesian information criterion (BIC) as an outcome measure. BIC distributions under varying polynomial order and number of clusters are shown in Fig. 4b, indicating a preference for an order-1 polynomial, and showing a slight improvement in BIC for three clusters compared to two clusters.

We then proceeded to fit a mixture model with three clusters and a polynomial order of 1 on the data. For this fitted model, we compared the distribution of LIR, MIR, and healthy classes across found clusters by defining cluster membership using the maximum membership probabilities (Fig. 4c), showing a total separation for LIR and healthy classes. For the MIR group, we see that 23 of the simulated trajectories are clustered separately from the LIR and healthy groups, while 2 trajectories are clustered with the healthy group.

When visualizing the predicted parameter trajectories using the mixture model, it can be seen that the differences between clusters that were encoded in the data generation are reconstructed by the mixture model. Fig. 4d-e show an increase in *k*_1_ and *k*_6_, coupled with a slight decrease in *k*_5_ for cluster 3 (LIR group), as well as a constant *k*_1_ and *k*_6_ with a strongly decreasing *k*_5_ for cluster 2 (MIR group).

### Mixture models accurately and efficiently categorize sparsely sampled unseen data

To assess the mixture model performance in a more uncertain situation, we simulated additional data in all three groups. However, while all simulated individuals had a measurement at *τ* = 0, between 3 and 5 additional measurements were taken at random intervals between 1 and 30 years, differing for each individual. As the other models were already not able to accomodate the high-quality sampled data, we only tested this experiment on the mixture model. Model parameters were estimated and the following trajectories were classified using the previously trained mixture model.

Fig. 5 shows the overall classification performance on the unseen data by the trained mixture model. Individuals belonging to the LIR group were all classified correctly, while the MIR and healthy groups had some minor overlap. Comparing overall performance by number of long time scale samples showed no strong performance decline with reduced time points.

**Fig 5.**
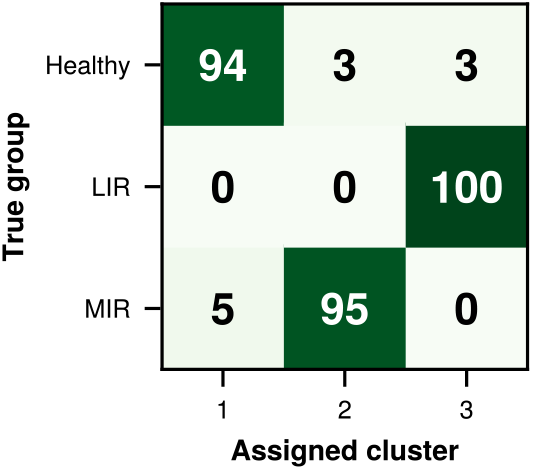
Mixture model performance on unseen data. Confusion matrix comparing the assigned labels with clusters assigned by the mixture model. The total number of individuals per group is 100.

## Discussion

In this work, we presented a novel multi-scale modelling strategy, combining LCTA with ODE modelling, to produce a hybrid framework that can be used to recover distinct subgroups in long-term disease progression from repeated short-term measurements in an unsupervised way. We have developed the `TemporalMixtureModels.jl` package in Julia which can be used to fit both univariate and multivariate mixture models on time series data. In a simulation study, we have simulated disease progression from a healthy state towards either liver, or muscle insulin resistance phenotypes using the EDES model. Using the `TemporalMixtureModels.jl` package, we show that the simulated subgroups can be reliably recovered from data. Furthermore, additional experiments with sparsely and irregularly sampled unseen data showed that a trained mixture model can accurately classify these individuals into the correct disease progression trajectories.

The approach is an extension of the Analysis of Dynamic Adaptations in Parameter Trajectories (ADAPT) work by van Riel et al. [13], where the evolution of a biological system over long-term time scale could be described by decoupling the short and long term dynamics in the system, and instead studying the trajectories of the the kinetic parameters estimated using data collected over the short-term time scale using splining approaches. While the study by van Riel, and also subsequent analyses by Tiemann et al [18] demonstrated the ADAPT approach could infer the expect parameter trajectories for unobserved state variables, the approaches assume one homogeneous population. Here we extend the approach to account for group-level heterogeneity in underlying parameter trajectories, which better reflects the heterogeneity we see in human populations.

We compared the mixture model to the approach described by Regazzoni et al. [14], where a neural ODE with a latent conditional parameter was trained to recover long term mechanistic model parameter trajectories. A large challenge with this approach is the assumption that all variability can be compressed down into a single conditional latent variable. In this work, we both reproduce the results from Regazzoni et al. on their windkessel model simulation, and we show that the neural ODE approach has great difficulty with representing the multivariate heterogeneity in the simulated data for the insulin resistance development case.

As neural networks are universal approximators, [26] this compression of multiple sources of variability into a single variable can technically be made within the training data, giving up potential extrapolation accuracy. However, this would substantially reduce the interpretability of the conditional latent variable. To recover interpretability, techniques such as symbolic regression [27] could potentially be used to recover the underlying equations. Alternatively, one could consider the use of multiple latent variables to circumvent this problem. However, the nonlinear nature of neural networks leaves no guarantee that the individual conditional latent variables describe separate sources of heterogeneity, and as a consequence they may be heavily correlated.

Specifically in human disease, the ability to properly handle multivariate groups is especially important, as progressive disease does not develop in isolation, but associate with many comorbidities. [28] Additionally, heavy crosstalk in human metabolism increases the complexity of disease progression. [29] Furthermore, in insulin resistance phenotypes, beyond glucose and insulin, many other metabolites, such a proteins [30], follow distinct postprandial profiles, further emphasizing the need for methods that can account for multivariate groupings in data.

An area where conditional neural networks could prove superior to the mixture models approach is in cases where the underlying heterogeneity contains a full spectrum of responses, rather than distinguishable patterns or subgroups. However, the complete grid search hyperparameters of `TemporalMixtureModels.jl` to analyze parameter trajectories ran in under 5 minutes, whereas the neural ODE approach took several hours to run in a single hyperparameter configuration on a desktop computer with an Intel Core i7-9700 processor. Therefore, even in a broad spectrum of responses, mixture models can serve as a good first step before trying neural network-based trajectory reconstruction methods (S6 Fig, S7 Fig). Additionally, when relying on data-driven methodology, such as neural networks and mixture models, the mixture model’s recovery of groups may be more favorable in biological settings, as these can be interpreted more easily a posteriori. Alternatively, in controlled experimental settings the ADAPT method [13] can effectively reconstruct parameter trajectories where underlying heterogeneity is not expected.

However, even in situations where a continuous spectrum of responses is observed, the use of a group-level distinction can aid in treatment selection for precision medicine [31]. Furthermore, the probabilistic nature of the grouping provided by mixture models still allows for precision treatment selection. [32, 33] Within type 2 diabetes, the characterization of individual phenotypes and associating these with specific treatments has shown to be effective for the design of dietary interventions, [34] showing that a method that can characterize disease progression trajectories and subsequently assign individuals into these categories with low data requirements is beneficial in precision medicine.

A limitation of this work is that we have only focused on simulated data to test our methodology, as prospective cohort studies with repeated OGTT experiments in the same individuals over several individuals are rare. However, we have attempted to simulate realistic data using a previously developed mechanistic model of OGTT response. Noise levels additionally resemble realistic values expected in glucose and insulin measurements, making the simulated test case relatively challenging.

Furthermore, the distinction between muscle and liver insulin resistance has been observed in real data. [25, 30] Nevertheless, while the model used for simulation contained specific parameters for liver gluconeogenesis and liver insulin response, the weak sensitivities of these parameters made them poor candidates for modulation to achieve a full liver insulin resistance phenotype. This highlights a challenge with simulating data in general, as the realism of the data simulated is limited by the realism of the model used to simulate it. In this work, we mitigated the limited sensitivity of the liver parameters by using gut absorption rate as a proxy for liver insulin sensitivity, resulting in glucose excursions comparable to real liver insulin resistance phenotypes. [30]

In this work, we show the mixture model on two use cases of progressive life-style related diseases. Progression of tissue specific insulin resistance, as well as progressive hypertension development. In this way, we demonstrate that the same framework can work for both cases. This strategy could be further applied to different multi-timescale problems in the future.

ODE models and mixture models have been combined before on equal time-scales, serving as an intermediate between population-based parameter estimation and full individualization of parameter estimates. [35, 36] However, the use of mixture models to describe temporal dynamics of model parameters presented here has not shown before. In the development of digital twins, this type of mutli-scale hybrid model would be effective when characterizing both the individual’s current disease state, as well as the most likely disease development trajectory. This information can be relevant in determining the proper treatment, and in assessing whether the given treatment has been effective. Furthermore, the key advantage of the mixture model is that it can be used with limited data, which increases its usability in clinical and precision medicine settings. Additionally, in future work, through the addition of a hidden Markov model, the framework presented here could be extended to allow for switching behaviour between trajectories, improving the representation of treatment effectivity. [37]

## Conclusion

By combining latent class trajectory analysis with ODE parameter estimation across time scales, we have introduced a novel modelling framework for multi-scale systems biology. We have shown the application of the framework in long-term parameter evolution simulation studies, and compared our strategy to a neural ODE-based strategy presented before, as well as a conditional neural network strategy. We have shown that under heavy sparsity, the mixture model framework provides a strong and stable grouping of individual trajectories. Furthermore, given a mixture model that is trained on a small-scale high quality dataset, the resulting model could be applied for reliable inference on sparsely and irregularly sampled data with high accuracy. Finally, we have released the developed methodology as a user-friendly Julia package that can be used and extended with a range of models beyond ODE models.

## Data and code availability

The code to reproduce the results in this manuscript is included in the GitHub repository https://github.com/ImmunoSystemsBiology/long-short-term. Source code for the TemporalMixtureModels.jl package can be accessed in the package GitHub repository https://github.com/Computational-Biology-TUe/TemporalMixtureModels.jl.

## Supporting information

**S1 Fig.**
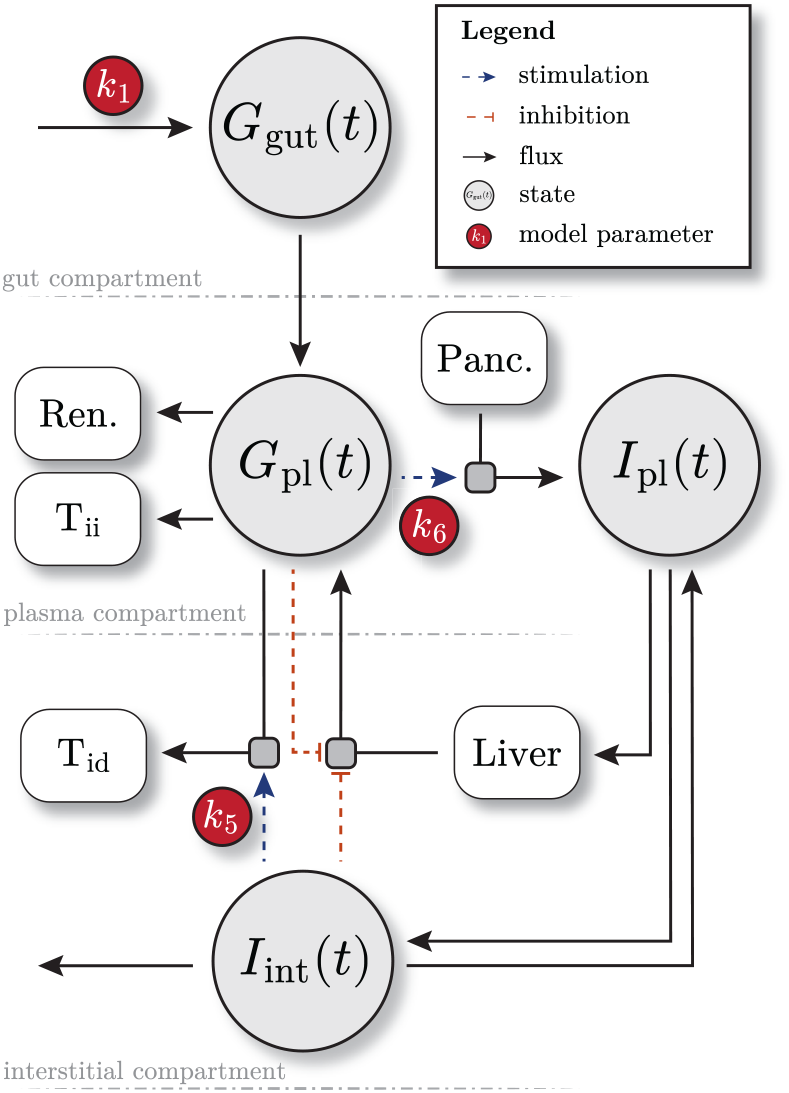
Illustration of the Eindhoven Diabetes Education Simulator (EDES). This figure shows the four compartments, fluxes, parameters, stimulations and inhibitions that are included in the EDES model. *G*_gut_(*t*): glucose in the gut compartment, *G*_pl_(*t*): glucose in the plasma compartment, *I*_pl_(*t*): insulin in the plasma compartment, *I*_int_(*t*): insulin in the interstitial compartment, Ren.: renal excretion of glucose, T_ii_: insulin-independent tissue glucose utilisation, T_id_: insulin-dependent tissue glucose utilisation, Panc.: pancreatic production of insulin.

### S2 Text. Eindhoven Diabetes Education Simulator model and parameter values

The Eindhoven Diabetes Education Simulator (EDES) model version used in this work consists of four state variables as a function of time *t*, depicting glucose in the gut (*G*_gut_(*t*)) and the plasma (*G*_pl_(*t*)), and insulin in the plasma (*I*_pl_(*t*)) and the interstitial compartment (*I*_int_(*t*)).

*Glucose in the gut* (in mg) is described by the following equation:

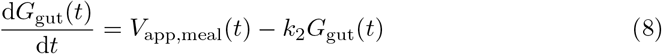

Where *k*_2_ describes the rate of glucose absorption from the gut into the plasma, and *V*_app,meal_(*t*) describes the glucose appearance from a meal into the gut compartment as:

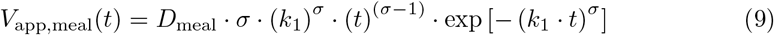

With shape parameter *σ* and rate of appearance parameter *k*_1_. *D*_meal_ is the meal glucose mass in mg.

*Glucose in the plasma* (in mmol *·* L^*−*1^) has the following equation:

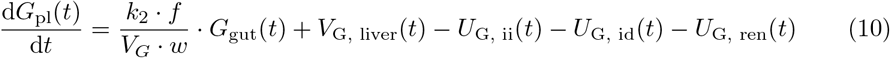

Where *f* is the conversion factor between glucose from mg to mmol, *V*_*G*_ is the glucose distribution volume in L per kg body weight and *w* is the individual’s body weight in kg.

The hepatic glucose flux *V*_G, liver_(*t*) is described by:

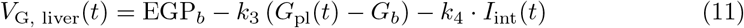

Where EGP_*b*_ is the fasting endogenous glucose production by the liver, *G*_*b*_ is the fasting glucose value, and *k*_3_ and *k*_4_ control the glucose and insulin effects on hepatic glucose production, respectively.

Insulin-independent glucose utilisation *U*_G, ii_(*t*) is described as:

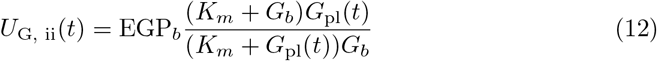

Where *K*_*m*_ is the Michaelis-Menten constant of glucose utilisation. Insulin-dependent glucose utilisation is described by:

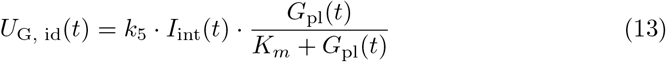

Where *k*_5_ represents the tissue insulin-sensitivity.

Finally, renal excretion of glucose *U*_G, ren_(*t*) has the equation:

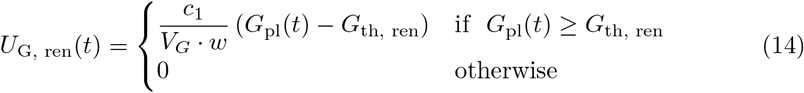

Where *c*_1_ is the renal excretion constant and *G*_th, ren_ is the plasma glucose concentration threshold above which renal excretion of glucose occurs.

*Insulin in the plasma* (in *µ*IU *·* mL^*−*1^) is described by the following equation:

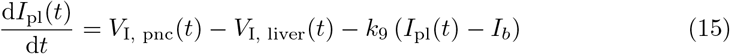

Where *k*_9_ represents the equilibrium constant between insulin in the plasma and interstitial insulin and *I*_*b*_ is the fasting insulin concentration.

Pancreatic production of insulin *V*_I, pnc_(*t*) is represented by the following equation:

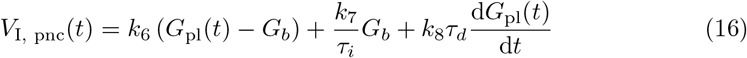

Where *k*_6_, *k*_7_ and *k*_8_ are the proportional, basal, and derivative rate parameters of glucose-stimulated insulin production. *τ*_*i*_ and *τ*_*d*_ are the time constants for *k*_7_ and *k*_8_, respectively.

Hepatic extraction of insulin *V*_I, liver_(*t*) is described as:

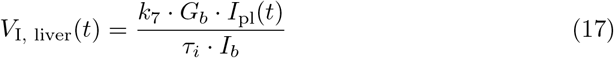

*Insulin in the interstitial space* is described by:

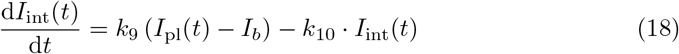

Where *k*_10_ describes the rate of insulin removal from the interstitial compartment.

Default parameter values, units, and their sources are included in S2.1 Table. Initial conditions for simulating the EDES model are shown in S2.2 Table.

**S2.1 Table.**
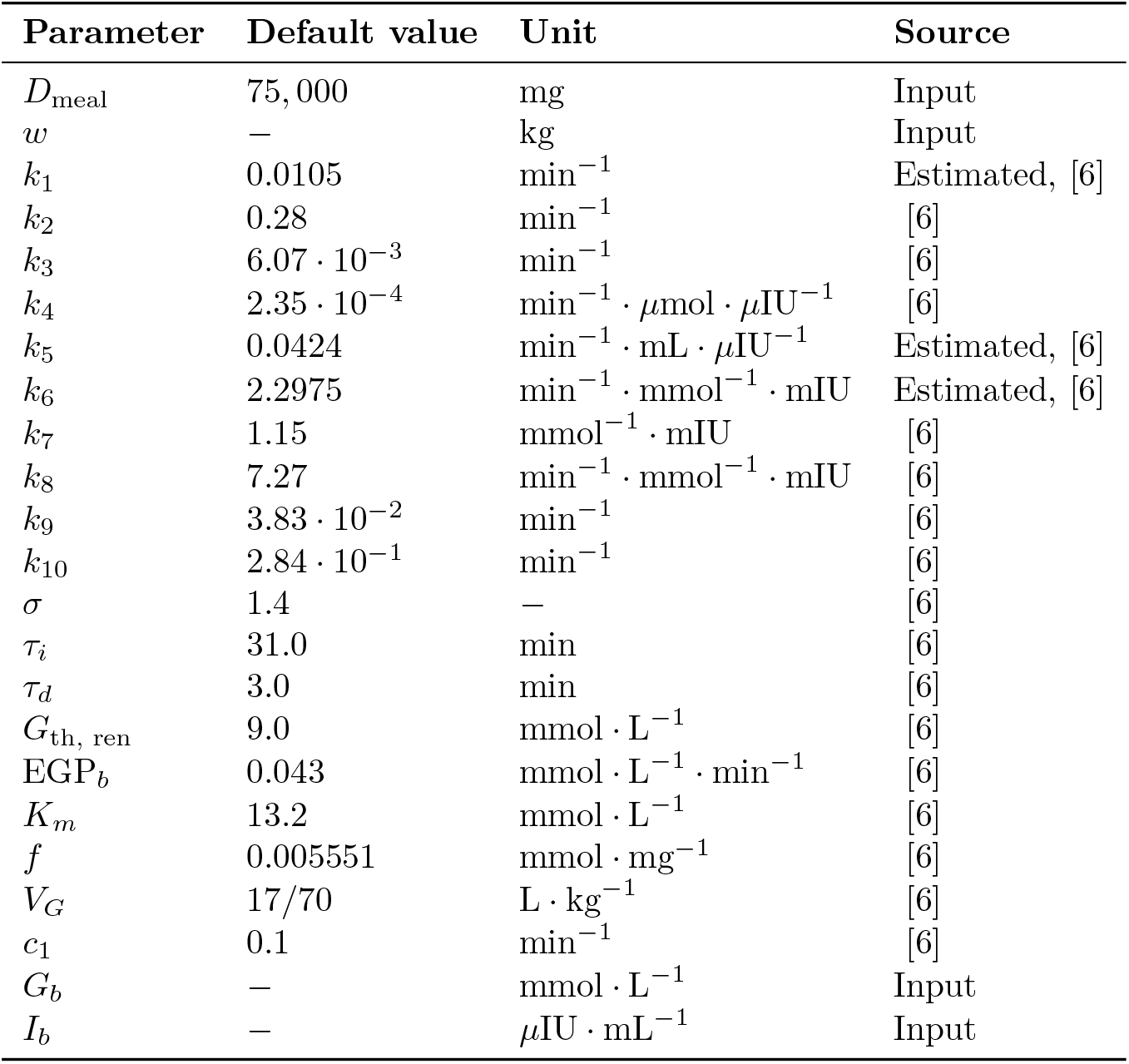
Parameter values for the EDES Model. This table includes the default parameter values, their units and their sources for all parameters in the EDES model. Parameters *D*_meal_, *w, G*_*b*_ and *I*_*b*_ need to be given as input to the simulation, and parameters *k*_1_, *k*_5_, and *k*_6_ are estimated from individual data. The remaining parameter values are left at their default values during parameter estimation.

| Parameter | Default value | Unit | Source |
|---|---|---|---|
| *D*<sub>meal</sub> | 75, 000 | mg | Input |
| *w* | − | kg | Input |
| *k*<sub>1</sub> | 0.0105 | min<sup>−1</sup> | Estimated, [6] |
| *k*<sub>2</sub> | 0.28 | min<sup>−1</sup> | [6] |
| *k*<sub>3</sub> | 6.07 · 10<sup>−3</sup> | min<sup>−1</sup> | [6] |
| *k*<sub>4</sub> | 2.35 · 10<sup>−4</sup> | min<sup>−1</sup> · µmol · µIU<sup>−1</sup> | [6] |
| *k*<sub>5</sub> | 0.0424 | min<sup>−1</sup> · mL · µIU<sup>−1</sup> | Estimated, [6] |
| *k*<sub>6</sub> | 2.2975 | min<sup>−1</sup> · mmol<sup>−1</sup> · mIU | Estimated, [6] |
| *k*<sub>7</sub> | 1.15 | mmol<sup>−1</sup> · mIU | [6] |
| *k*<sub>8</sub> | 7.27 | min<sup>−1</sup> · mmol<sup>−1</sup> · mIU | [6] |
| *k*<sub>9</sub> | 3.83 · 10<sup>−2</sup> | min<sup>−1</sup> | [6] |
| *k*<sub>10</sub> | 2.84 · 10<sup>−1</sup> | min<sup>−1</sup> | [6] |
| *σ* | 1.4 | − | [6] |
| *τ*<sub>*i*</sub> | 31.0 | min | [6] |
| *τ*<sub>*d*</sub> | 3.0 | min | [6] |
| *G*<sub>th, ren</sub> | 9.0 | mmol · L<sup>−1</sup> | [6] |
| EGP<sub>*b*</sub> | 0.043 | mmol · L<sup>−1</sup> · min<sup>−1</sup> | [6] |
| *K*<sub>*m*</sub> | 13.2 | mmol · L<sup>−1</sup> | [6] |
| *f* | 0.005551 | mmol · mg<sup>−1</sup> | [6] |
| *V*<sub>*G*</sub> | 17/70 | L · kg<sup>−1</sup> | [6] |
| *c*<sub>1</sub> | 0.1 | min<sup>−1</sup> | [6] |
| *G*<sub>*b*</sub> | − | mmol · L<sup>−1</sup> | Input |
| *I*<sub>*b*</sub> | − | µIU · mL<sup>−1</sup> | Input |

**S2.2 Table.** Initial conditions for the EDES model. This table includes the used initial conditions for simulating the EDES model.

| **State** | **Initial condition** | **Unit** |
|---|---|---|
| *G*<sub>gut</sub>(*t*) | 0 | mg |
| *G*<sub>pl</sub>(*t*) | *G*<sub>*b*</sub> | mmol · L<sup>−1</sup> |
| *I*<sub>pl</sub>(*t*) | *I*<sub>*b*</sub> | *μ*IU · mL<sup>−1</sup> |
| *I*<sub>int</sub>(*t*) | 0 | *μ*IU · mL<sup>−1</sup> |

**S3 Table.**
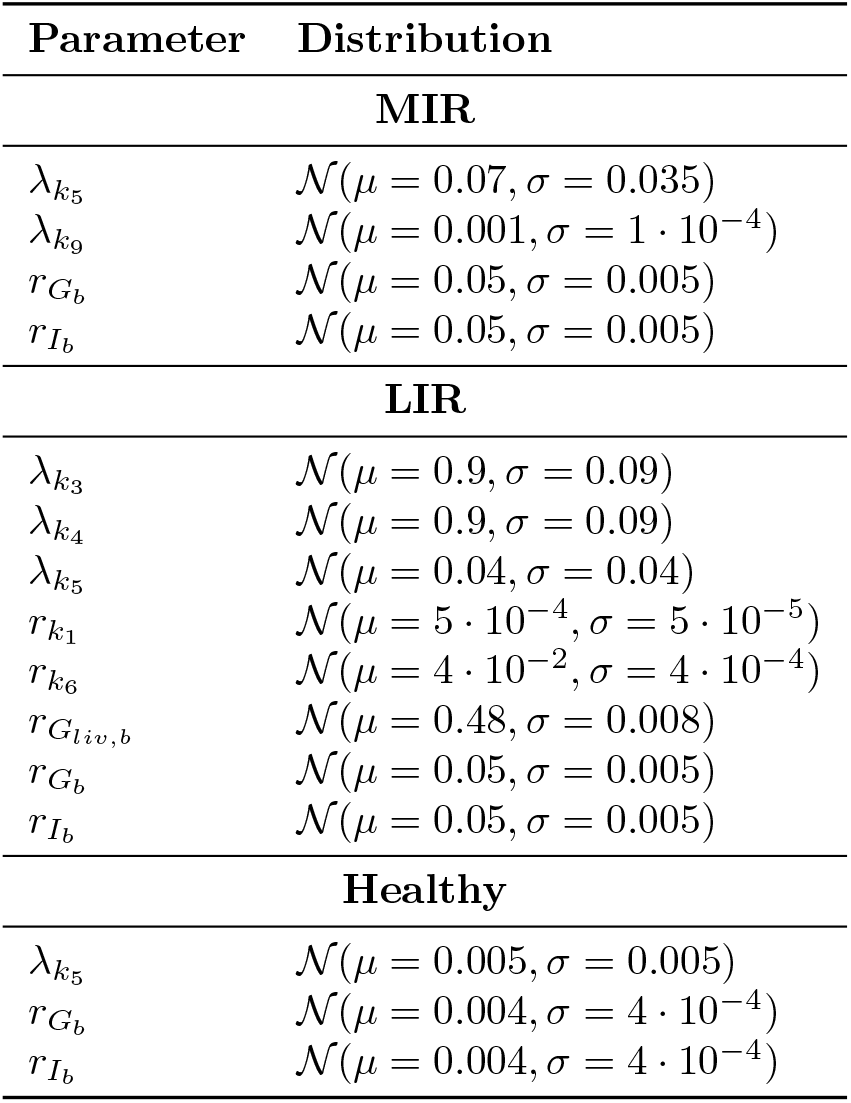
Progression model parameter values for virtual populations. Decay (*λ*) and rate (*r*) parameters for muscle insulin resistance (MIR), liver insulin resistance (LIR) and healthy groups. Samples were clamped to zero on the lower end to prevent negative parameter values.

| **Parameter** | **Distribution** |
|---|---|
| **MIR** |  |
| λ<sub>k<sub>5</sub></sub> | 𝒩(μ = 0.07, σ = 0.035) |
| λ<sub>k<sub>9</sub></sub> | 𝒩(μ = 0.001, σ = 1 · 10<sup>−4</sup>) |
| r<sub>G<sub>b</sub></sub> | 𝒩(μ = 0.05, σ = 0.005) |
| r<sub>I<sub>b</sub></sub> | 𝒩(μ = 0.05, σ = 0.005) |
| **LIR** |  |
| λ<sub>k<sub>3</sub></sub> | 𝒩(μ = 0.9, σ = 0.09) |
| λ<sub>k<sub>4</sub></sub> | 𝒩(μ = 0.9, σ = 0.09) |
| λ<sub>k<sub>5</sub></sub> | 𝒩(μ = 0.04, σ = 0.04) |
| r<sub>k<sub>1</sub></sub> | 𝒩(μ = 5 · 10<sup>−4</sup>, σ = 5 · 10<sup>−5</sup>) |
| r<sub>k<sub>6</sub></sub> | 𝒩(μ = 4 · 10<sup>−2</sup>, σ = 4 · 10<sup>−4</sup>) |
| r<sub>G<sub>liv,b</sub></sub> | 𝒩(μ = 0.48, σ = 0.008) |
| r<sub>G<sub>b</sub></sub> | 𝒩(μ = 0.05, σ = 0.005) |
| r<sub>I<sub>b</sub></sub> | 𝒩(μ = 0.05, σ = 0.005) |
| **Healthy** |  |
| λ<sub>k<sub>5</sub></sub> | 𝒩(μ = 0.005, σ = 0.005) |
| r<sub>G<sub>b</sub></sub> | 𝒩(μ = 0.004, σ = 4 · 10<sup>−4</sup>) |
| r<sub>I<sub>b</sub></sub> | 𝒩(μ = 0.004, σ = 4 · 10<sup>−4</sup>) |

**S4 Fig.**
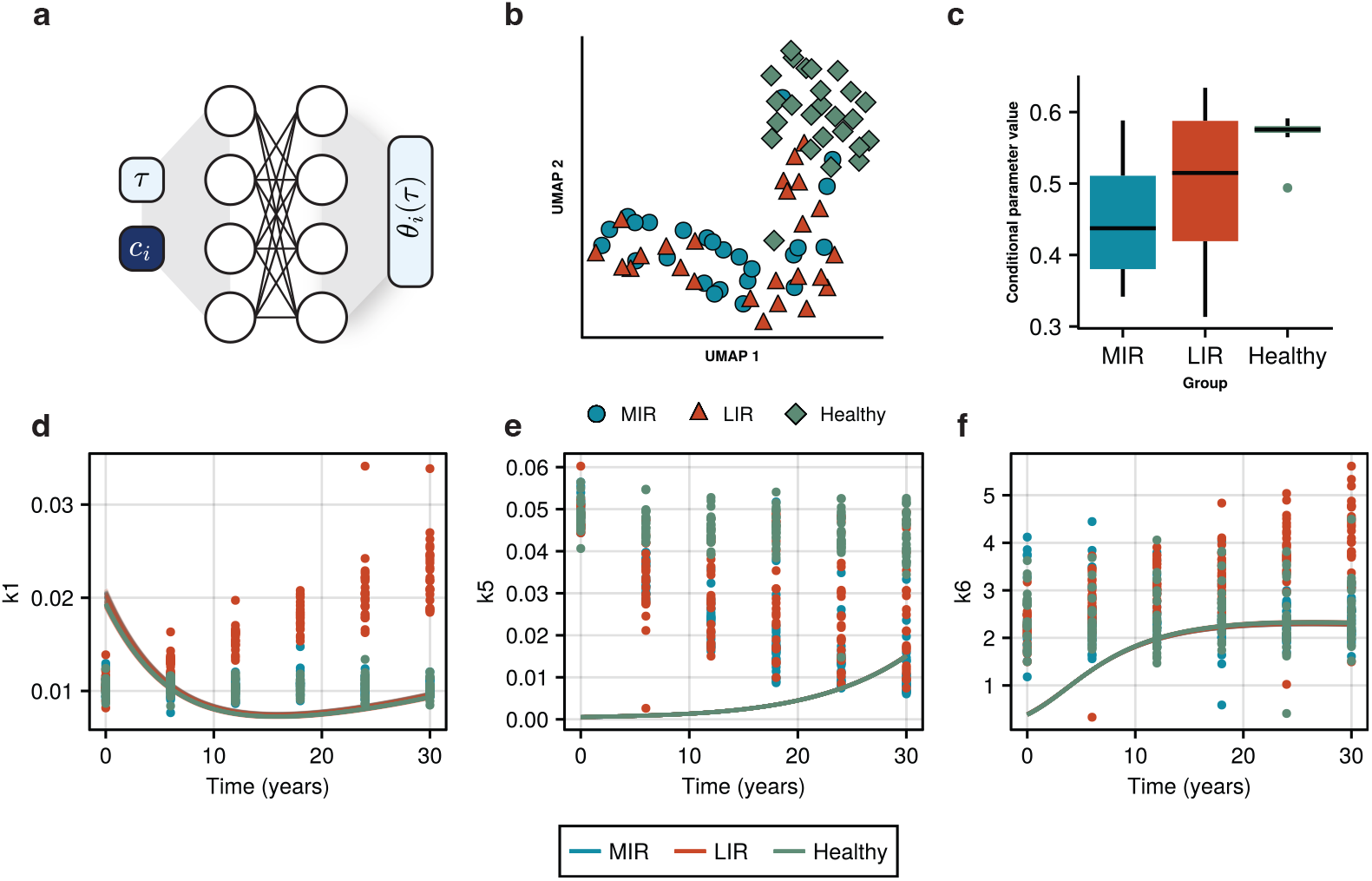
Conditional neural network reconstruction of parameter trajectories. Results of the reconstruction of the parameter trajectories of the MIR, LIR, and healthy subgroups using a conditional neural network. A: Illustration of the structure of the conditional neural network, where a neural network is used to describe the raw parameter values as a function of time (*τ* ) and a conditional parameter. The conditional parameter *c*_*i*_ is constant over time and allows for modelling specific interindividual differences. B: UMAP plot of the trained conditional parameter across 25 optimization runs with different initial parameters, colored by subgroup (blue circle: MIR, orange triangle: LIR, green diamond: healthy). C: Representative boxplot of the conditional parameter values across different groups for a single optimization run. D-F: Reconstructions of the parameter trajectories (lines) for the parameters *k*_1_, *k*_5_, and *k*_6_, respectively. Lines and ground truth parameter values are colored by subgroup.

### S5 Text. Two-element Windkessel Model Experiment

To verify the applicability of the mixture model approach in a different situation, we replicated the simulation described by Regazzoni et al. [14], describing the varying hypertension developments in a two-element windkessel model of blood circulation.

The two-element windkessel model describes the peripheral and distal blood pressure (*p*_*p*_ and *p*_*d*_) through peripheral and distal resistances (*R*_*p*_ and *R*_*d*_) and compliances (*C*_*p*_ and *C*_*d*_), in a simple hydrodynamic setup. The ordinary differential equations for this model are:

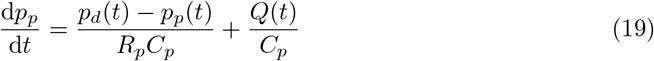

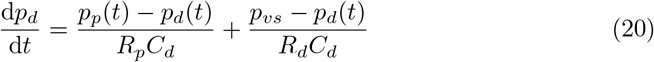

The development of hypertension was then simulated through a set of differential equations describing the change of the distal resistance and compliance over the time course of weeks, governed by a severity parameter *α ∈* (0, 1):

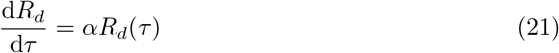

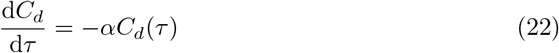

Data for 100 virtual patients was simulated by randomly sampling a combination of parameter values for *C*_*p*_, *C*_*d*_, *R*_*p*_, *R*_*d*_, and *α*, within the bounds shown in S5.1 Table and simulating equations 19 and 20 for 20 heartbeats. Where available, parameter values from Regazzoni et al. were used.

**S5.1 Table.** Parameter bounds for the two-element Windkessel model. This table includes the minimum and maximum values for parameters selected for the simulation of the two-element Windkessel model.

| **Parameter** | **Minimum** | **Maximum** |
|---|---|---|
| *R<sub>p</sub>* | 8.1757 · 10<sup>−2</sup> | 1.7326 · 10<sup>−1</sup> |
| *C<sub>p</sub>* | 3.1197 · 10<sup>−2</sup> | 3.1464 · 10<sup>−2</sup> |
| *R<sub>d</sub>* | 6.9006 · 10<sup>−1</sup> | 1.1599 |
| *C<sub>d</sub>* | 1.1851 | 2.1731 |
| *α* | 0.0 | 1.0 · 10<sup>−2</sup> |

As *p*_*vs*_ and *Q*(*t*) were unavailable, a value of *p*_*vs*_ = 5mm Hg was selected as used in [38]. For *Q*(*t*), a simple sinusoidal wave was selected:

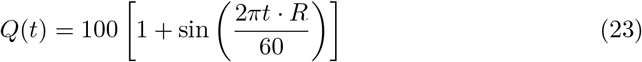

With *R* representing the heart rate, which was set at 75 bpm. For simulating measurements, the first 10 heartbeats were discarded, to allow the model to find its stationary state. Only *p*_*p*_(*t*) was measured at intervals Δ*t* = 1 *·* 10^*−*2^ s, and additive measurement error was added using a normally distributed error model with standard deviation *ρ* = 0.75 mm Hg^1*/*2^.

Subsequently, model simulations were repeated while simulating the long-term model using equations 21 and 22, taking measurements every week from week 0 until week 50.

From the simulated data, all four model parameters, the initial condition for *p*_*d*_, and the standard deviation of the residual for *p*_*p*_ (*σ*_*p*_) were estimated for each individual and each week separately. From the resulting trajectories, 25 virtual patients were selected randomly as a train set. A polynomial mixture model was trained on the trajectories of these 25 patients for parameters *R*_*d*_ and *C*_*d*_ with polynomial order 1 and *K* = 3 components. The model was subsequently tested on the entire population of 100 virtual patients.

**S6 Fig.**
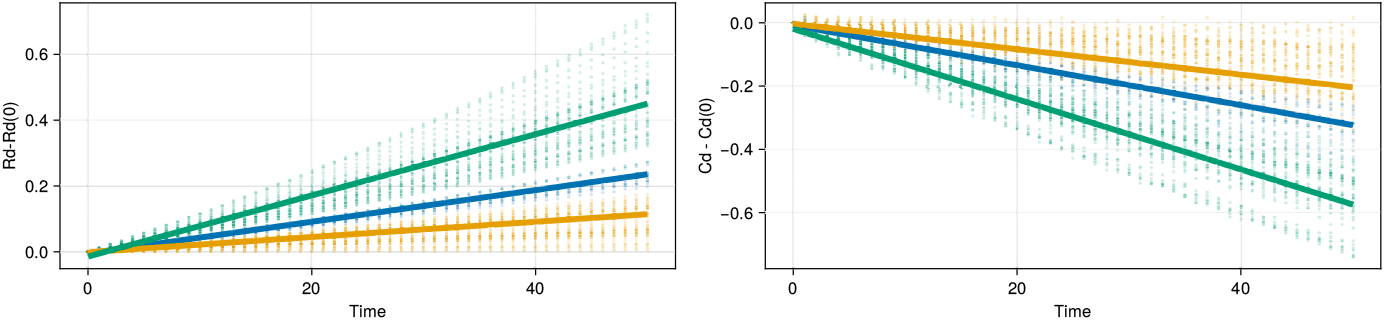
Group-level trajectories for a 3-component mixture model on the hypertension development simulation. Data was normalized to reflect relative change with respect to the initial parameter values. The mixture model clearly identified three groups within the data (green, blue, and orange lines), which reflect the progressions in the data (green, blue, and orange dots).

**S7 Fig .**
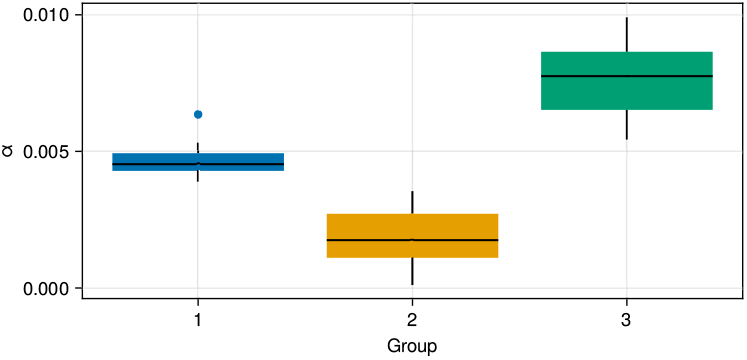
Distribution of alpha values per mixture model group. For each group in the mixture model, distributions of *α* values known from the simulation are shown.

**S8 Fig.**
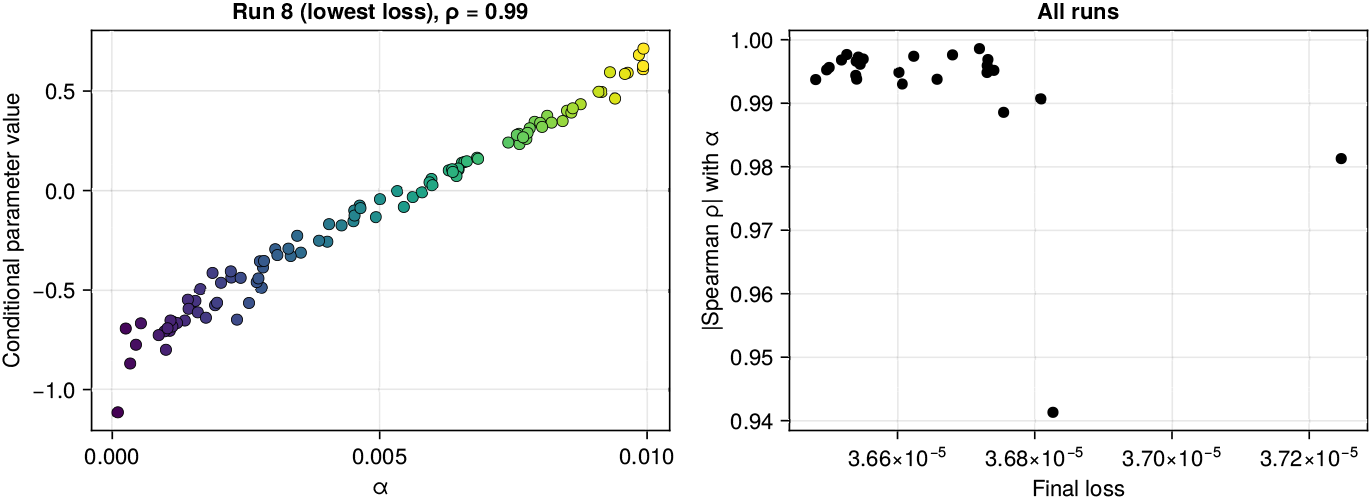
Correlation between the conditional parameter of the neural ODE and alpha. For each individual in the simulated windkessel data, the conditional neural ODE model was fit. The resulting conditional parameter reveals a strong correlation with the original *α* parameter for all 25 independent rounds of parameter estimation

**Fig S9.**
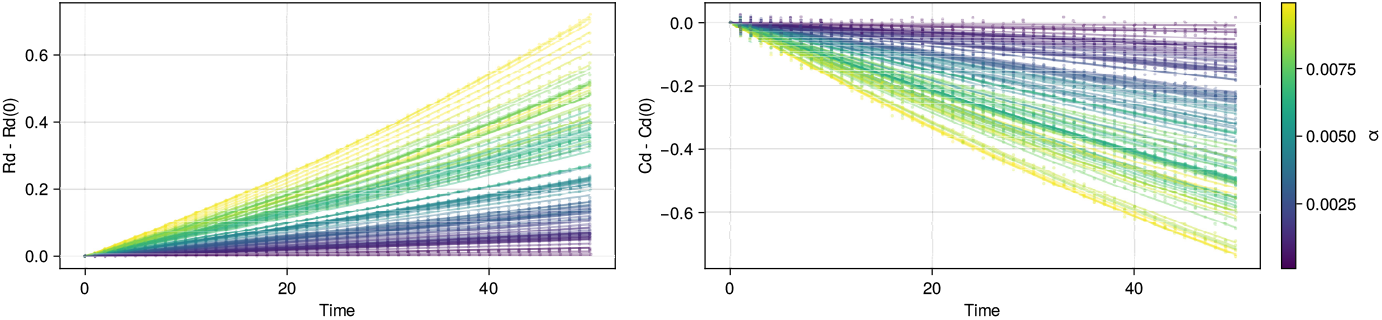
Simulated model behavior for different conditional parameter values in the neural ODE. Similarly to what was shown by Regazzoni et al. [14], the neural ODE model shows strong agreement with the original simulated data.

